# Fast Phenotype Simulation for Genotype Representation Graphs

**DOI:** 10.1101/2025.08.15.670378

**Authors:** Aditya Syam, Chris Adonizio, Xinzhu Wei

## Abstract

**Motivation:** The Genotype Representation Graph (GRG) [DeHaas et al., 2025] is a graph representation of whole genome polymorphisms, designed to encode the variant hard-call information in phased whole genomes. It encodes the genotypes as an extremely compact graph that can be traversed efficiently, enabling dynamic programming-style algorithms on applications such as genome-wide association studies that run faster on biobank-scale data than existing alternatives. To facilitate scalable statistical genetics, we present *GrgPhenoSim*, an extremely fast phenotype simulator for GRGs, suitable for simulating phenotypes on biobank-scale datasets.

**Results:** *GrgPhenoSim* contains all the primary functionalities of a phenotype simulator, uses a standardized output, and supports customized simulations. *GrgPhenoSim* is dozens to hundreds of times faster than *tstrait* [Tagami et al., 2024], *a fast ancestral recombination graph-based phenotype simulator, when the sample size ranges from thousands to hundreds of thousands samples*.

**Availability:** The GrgPhenoSim library and use-case demonstrations are available at https://github.com/aprilweilab/grg_pheno_sim

The documentation for GrgPhenoSim is hosted at https://grgl.readthedocs.io/en/latest/index.html

## 1 Introduction

Phenotypes, such as height (continuous) or type II diabetes (binary), refer to observable traits of an individual. Genome-wide association studies (GWAS), fine mapping, and heritability analysis are important for understanding the genetic architecture of phenotypes. Phenotype simulation is a pivotal part of these applications because it provides a ground truth for testing and evaluating new statistical methods. Causal variants are genetic variants that affect phenotypic values. Within the scope of phenotype simulation, causal variants are often sampled from genotype data, and then the phenotypes are computed based on the genotypes. Conventionally, such simulations use tabular genotype data structures such as the variant call format (VCF) [Danecek et al., 2011] and the PLINK [Chang et al., 2015] BED file format as input. However, as genotype datasets are becoming larger and larger, for example, storing 200,000 phase UK Biobank wholegenome polymorphisms in VCF.gz (gzip compressed VCF files) requires 2.13 terabytes, it becomes increasingly expensive to perform repeated phenotype simulations on tabular genotype data [DeHaas et al., 2025].

An ancestral recombination graph (ARG) is a data structure that provides an efficient summary of the ancestry of population-level genetic data through coalescence and recombination events [Nielsen et al., 2025]. ARGs can compactly store local genealogies along the genome that show how genetic materials have been recombined and inherited over time. Some ARG data formats, such as the tree sequence in tskit format[Kelleher et al., 2018], can also represent genotype data. The Python library tstrait [Tagami et al., 2024], building on this format, enables more efficient simulation of phenotypes than simulations based on tabular genotype data. However, ARG inference is currently infeasible (or extremely expensive) for biobank-scale whole-genome sequencing (WGS) datasets, hence limiting the utility of tstrait [Tagami et al., 2024] on real datasets.

Here, we leverage the genotype representation graph (GRG) [DeHaas et al., 2025] as the input to perform graph-based phenotype simulation. A GRG is a multi-tree structure that losslessly encodes the hard-call information in phased genotypes. The graph hierarchy efficiently represents information without using any compression library. Constructing GRG files for all autosomes of the 200,000 UK biobank WGS dataset cost a total of 80 pounds on the UK Biobank DNANexus cloud computing platform, yielding an uncompressed GRG that is 13 times smaller than the same dataset in compressed VCF (VCF.gz) format [DeHaas et al., 2025]. Moreover, the dot product calculation via traversing over a GRG runs faster than all tested alternatives (PLINK [Chang et al., 2015], XSI [Wertenbroek et al., 2022], and Savvy [LeFaive et al., 2021]) and enabled extremely efficient GWAS [DeHaas et al., 2025]. Here, we leverage GRG and its dot product function to build a highly efficient phenotype simulator to facilitate statistical method development. Moreover, we demonstrate the use of linear algebra operations on the standardized genotype matrix (an implicit transformation of the GRG), showing that the GRG dot product can be leveraged to speed up other statistical and population genetics calculations beyond phenotype simulation.

## 2 Methods

### 2.1 Model

#### 2.1.1 A Brief Overview of GRG Dot Product

A GRG [DeHaas et al., 2025] is a directed acyclic graph with nodes that contain zero or more “mutations”. In real data, the mutation status is often unknown, in which case either the derived allele or the minor allele can be used as “mutation”. The sample nodes correspond to the leaf nodes of the graph, which have no successors. Each haploid genome is represented as one sample node and each diploid genome is represented as two sample nodes. A path from the node containing mutation m_i_ to the sample node s_j_ exists if and only if the genotype for s_j_ contains mutation m_i_, therefore encoding the same information as tabular genotype hard calls. A GRG [DeHaas et al., 2025] has down edges that are directed from the roots to the sample nodes (Fig. 1), and these edges carry forward the mutations down the tree.

**Figure 1:**
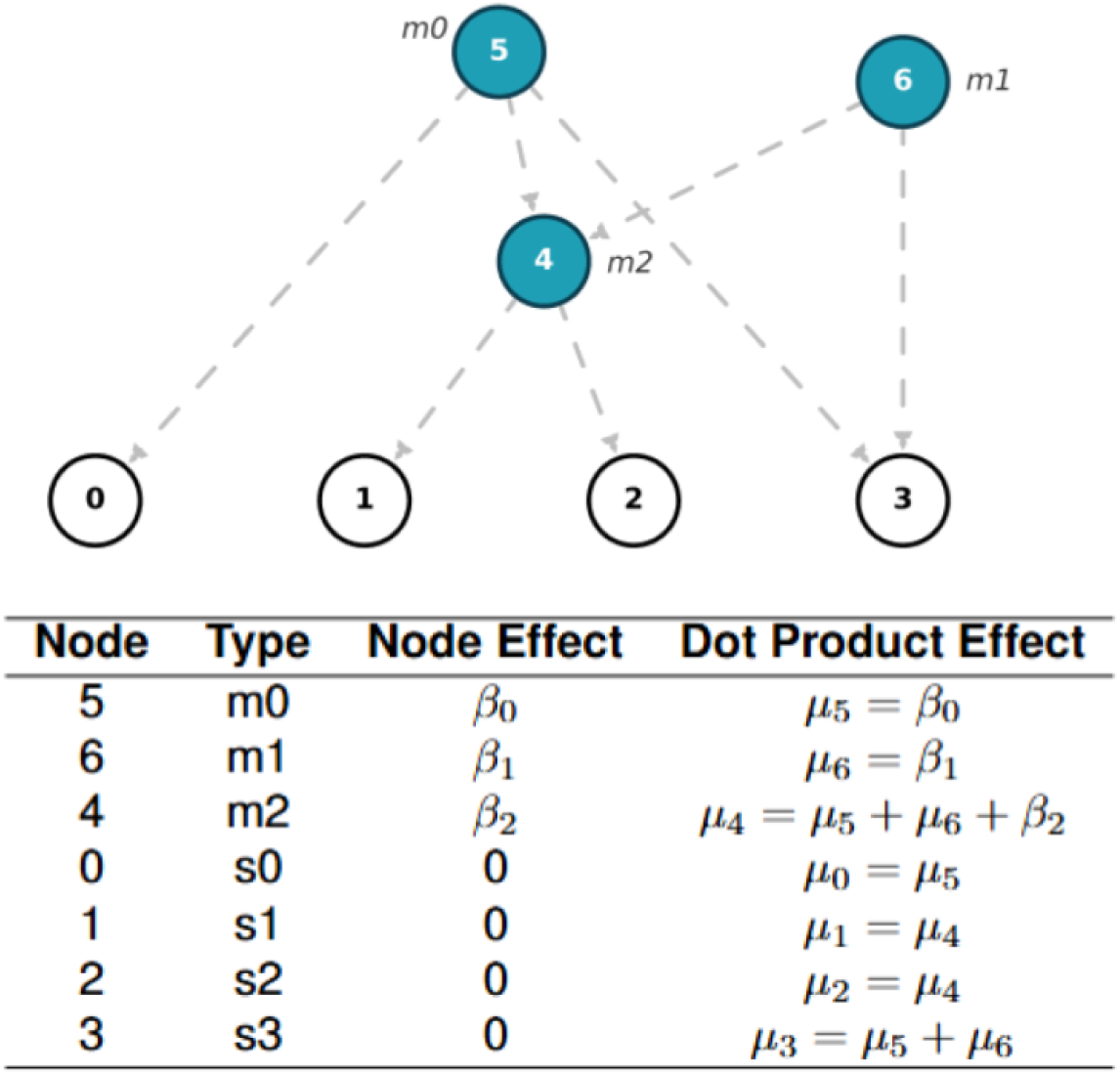
A demonstration of GRG traversal for computing dot product values for phenotype simulation. The upper image shows a GRG, with node identifiers labeled within each node, and mutation identifiers labeled alongside their associated nodes. Every node in the GRG corresponds to a unique set of samples a node can reach, with individual mutations being associated with specific nodes. For instance, node 4 has two children - nodes 1 and 2, meaning that m2 (associated with node 4) is carried by nodes 1 and 2, but not by nodes 0 and 3. Finally, pairs of haploid sample nodes combine to form a single diploid sample (node 0 and 1 form the genotype for a hypothetical individual 1). The table below shows the graph traversal order and the corresponding dot product value at each node. In the context of phenotype simulation, node effect (input to dot product) is simply the causal phenotypic effects from the associated mutations with that node. The dot product function computes the dot product effect at a node by adding the dot product effects from all its parent nodes with its own node effect. For the sample nodes 0,1,2,3, the dot product effects (function output) are the genetic values of these corresponding haploid genomes.

#### 2.1.2 Efficient Matrix-Vector Operations with GRG

As discussed, each node within the GRG is mapped to one or more mutations [DeHaas et al., 2025]. The mutations contained within the sample nodes are the cumulative effect of the mutations contained within all ancestral nodes. In order to compute genetic values for these samples, a graph traversal is necessary to ensure that the effect sizes are passed down through all ancestral nodes, right until the samples. The dot product operation in the GRG Library (GRGL) utilizes a highly optimized graph traversal method to compute the equivalent dot product result [DeHaas et al., 2025], such that matrix algebra operations can be easily implemented.

More specifically, let **G** be a haploid (phased) genotype matrix of size 2*N*-by-*M*, where *N* is the number of diploid individuals/genomes, and *M* is the number of variants. Each element in **G** takes a value 0 or 1. Traversals of the GRG that only sum the values calculated transitively for each node’s children (upward traversal) are equivalent to the matrix product **G**^**T**^**v**, where **v** can be any 2*N*-by-1 real vector. Similarly, traversals that only sum values calculated transitively for parents (downward traversal) are equivalent to the matrix product **Gu**, where **u** can be any *M* -by-1 real vector (1). With the newly updated GRGL Python API v2.2, both types of dot products can be done efficiently via the function pygrgl.dot product() [Wei Lab, 2025].

#### 2.1.3 Computing the Additive Genetic Value

Let **X** be a diploid genotype matrix of size *N* - by-*M*, such that each entry of **X** can take a value of 0, or 1, or 2. Dot products involving **X** or **X**^*T*^ could be done via GRG by identifying mathematically equivalent operations based on the dot products involving **G** or **G**^*T*^, because the additive (i.e., linear) relationship **X**_*i,j*_ = **G**_2*i*−1,*j*_ + **G**_2*i,j*_ holds at each entry with row index *i* column index *j*.

The standard equation for computing continuous phenotypes under an additive genetic model is: **y** = **X*β*** + ***ϵ***. Here, **y** is the *N*-by-1 phenotype vector, and ***ϵ*** designates the *N*-by-1 environmental noise vector. ***β*** is the *M* -by-1 effect size vector, where entries corresponding to the causal variants that have nonzero causal effects and entries corresponding to non-causal variants are set to zero. We then call the pygrgl.dot product() function to compute **G*β*** to get the additive genetic value for each haploid genome (Fig. 1). Adding the genetic values at the (2*i* − 1)-th and 2*i*-th haploid genomes together, we get the *i*-th diploid individual’s genetic value (Fig. 1).

#### 2.1.4 Enabling Standardized Genotype Matrix

Standardized genotype matrices are frequently used in statistical genetics, but phenotype simulators, such as GCTA [Yang et al., 2011] and *tstrait* [Tagami et al., 2024*], often do not provide such an option. In GrgPhenoSim*, we provide the option “standardized” for simulating phenotypes based on standardized diploid genotype matrix 𝒳, where 𝒳 = (**X** − **U**)**Σ**. Here, **U** is an *N* - by-*M* matrix where each entry in the *i*-th column is 2*f*_*i*_, twice the frequency of the mutation allele *i*. **Σ** is a diagonal matrix where each *i*-th diagonal element is the inverse standard deviation at a mutation, 1*/σ*_*i*_, with 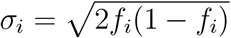.

Briefly, in order to efficiently compute the genetic value 𝒳***β***, we calculate **X**(**Σ*β***) − **UΣ*β***. Here, **Σ*β*** is a *M* -by-1 vector with elements *β*_*i*_*/σ*_*i*_ at each *i*-th entry. Therefore, **X**(**Σ*β***) can be computed via dot product between matrix **G** and vector **Σ*β***. After that, we perform an element-wise subtraction by ∑ _*i*_ 2*f*_*i*_*β*_*i*_*/σ*_*i*_,because **UΣ*β*** is simply a *N*-by-1 vector filled with the same scalar that takes the value ∑_*i*_ 2*f*_*i*_*β*_*i*_*/σ*_*i*_.

When the “standardized” option is evoked, the causal effect sizes are by default sampled from a normal distribution *β ∼* 𝒩 (0, *h*^2^*/M*_*causal*_), where *M*_*causal*_ is the number of causal mutations, and *h*^2^ is the narrow sense heritability. Then, after computing genetic value 𝒳***β***, environmental noise is sampled from *ϵ* ∼ 𝒩 (0, *V ar*(𝒳***β***)(1*/h*^2^ − 1)) to ensure that heritability is equal to *h*^2^.

### 2.2 Computational Pipeline

The simulation pipeline processes GRGs through four main steps to generate phenotype data. The implementation utilizes *numpy* [Harris et al., 2020] for numeric operations (enabling standard math libraries such as BLAS) and *pandas* [pandas development team, 2020] for data manipulation and output. The pipeline (Fig. 2) starts with simulating causal effects, then proceeds with computing genetic values, adding environmental noise, and normalizing phenotypic values. The first step involves obtaining effect sizes for mutations in the GRG. Users can either employ one of the built-in distributions or input custom effect sizes. The built-in statistical distributions include normal, exponential, fixed, gamma, and t-distributions, available for both univariate and multivariate (for jointly simulating multiple phenotypes) settings. When “standardized” genotype matrix is evoked, normal distribution will be used for simulating causal effect size. The dataframe contains the *mutation id, effect size*, and *causal mutation id*.

**Figure 2:**
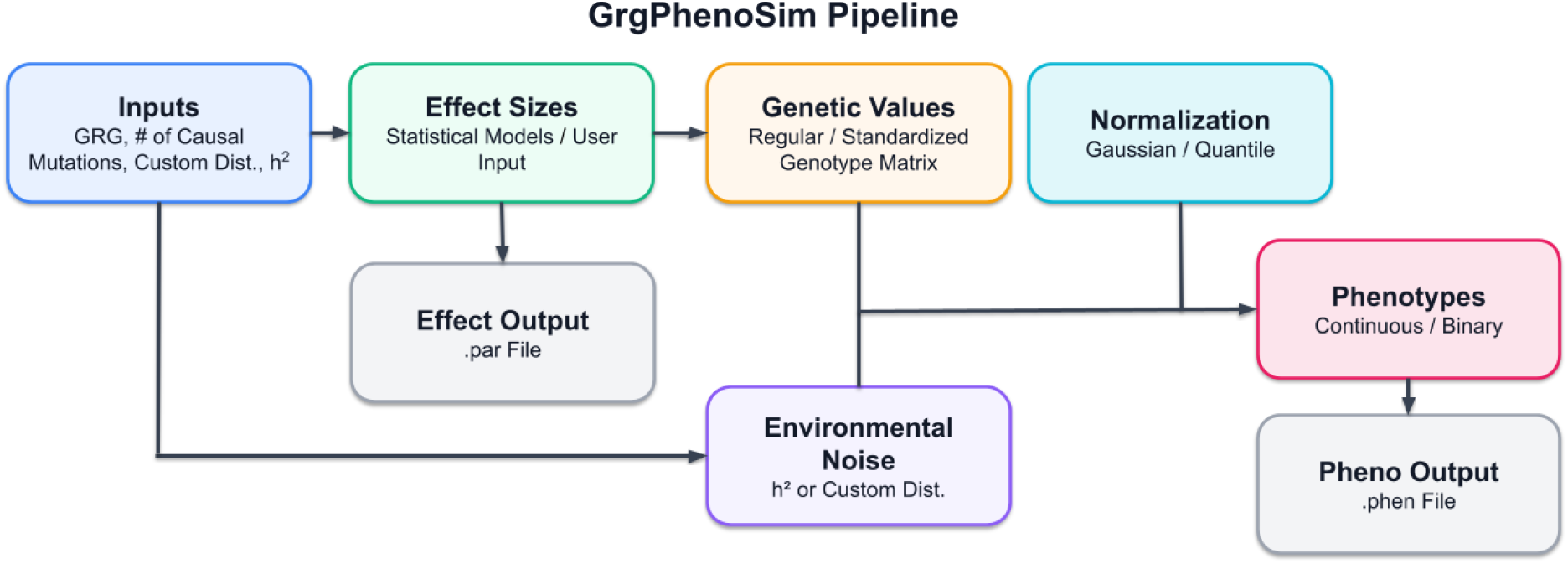
Visualization of the simulation pipeline.

The second stage is to compute genetic values for diploid genomes, leveraging GRG dot product function [DeHaas et al., 2025] (Fig. 1). The system generates dataframes first at the sample level (*sample node id, genetic value, causal mutation id*), and then at the individual level (*individual id, genetic value, causal mutation id*). Users may opt for normalizing genetic values at this stage.

The integration of environmental noise *ϵ* forms the third step, where the noise is generated through either an input narrow-sense heritability *h*^2^ or a user-defined distribution. If using an input *h*^2^, *ϵ* ∼ 𝒩 (0, *V ar*(𝒳***β***)(1*/h*^2^ − 1)) when “standardized” genotype matrix is evoked, otherwise *ϵ* ∼ 𝒩 (0, *V ar*(**X*β***)(1*/h*^2^ − 1)) with the original genotype matrix **X**. This stage produces a dataframe containing *causal mutation id* (for multiple causal mutations), *individual id, genetic value, environmental noise*, and *phenotype* values.

The final optional normalization step offers either standard normalization or quantile normalization. Users can normalize either phenotypes alone or both genetic and phenotype values. For quantile normalization, the output dataframe includes all previous columns plus *normalized phenotype*.

### 2.3 Additional Features

*GrgPhenoSim* supports binary phenotype simulation. This step follows the standard pipeline to simulate a standardized continuous phenotype, then employs a *population prevalence* parameter to establish a Gaussian threshold to convert continuous values into binary outcomes. Lastly, the system overwrites continuous phenotypes with their binary counterparts.

Further, *GrgPhenoSim* can accommodate multiple GRG inputs simultaneously, particularly useful when chromosome-specific genotype files are available, without having to merge them into a single file. This functionality requires consistent sample orders across GRGs and offers both parallel and sequential loading options. Genetic values from multiple GRG inputs are combined prior to noise simulation and final phenotype computation.

*GrgPhenoSim* can output standard .*par* files same as GCTA containing columns for *mutation id, AlternateAllele, Position, RefAllele, Frequency*, and *Effect* [Yang et al., 2011]. *It can also output standard* .*phen* files [Yang et al., 2011] containing *person id* and *phenotypes* columns, with customizable header options.

This comprehensive pipeline (Fig. 2)ensures efficient phenotype simulation while being highly adaptable to individual users’ needs.

### 2.4 Usage

*GrgPhenoSim* is implemented and wrapped in Python, and uses functionalities from the *GRG* library written in C++ to ensure efficiency. *GrgPhenoSim* offers two primary usage patterns: step-by-step execution through sequential notebooks, or a streamlined wrapper method for end-to-end simulation. A minimal example for continuous phenotype simulation is provided to showcase the simplicity of the Python interface.

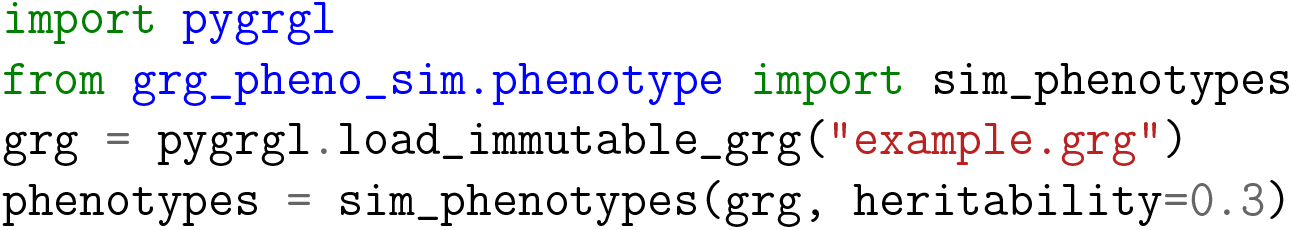

## 3 Results

### 3.1 Validation

Intermittent outputs were verified to ensure that the simulator works properly. For example, the GRG *dot product* function was verified by testing against a recursive algorithm based on GRG, and against matrix multiplication based on the original genotype matrix. The genetic values computed by *GrgPhenoSim* were also cross-verified against *tstrait* [Tagami et al., 2024]. *In this validation, simulated tree sequences were converted to GRGs using GRGL. Identical effect sizes were fed into both GrgPhenoSim* and *tstrait* [Tagami et al., 2024], *and the resulting genetic values (i*.*e*., *Xβ*) were identical.

### 3.2 Runtime Efficiency

We benchmarked the scalability of *GrgPhenoSim* against the current state-of-the-art graph-based phenotype simulator *tstrait* [Tagami et al., 2024], with respect to sample size and number of causal mutations (Fig. 3).

**Figure 3:**
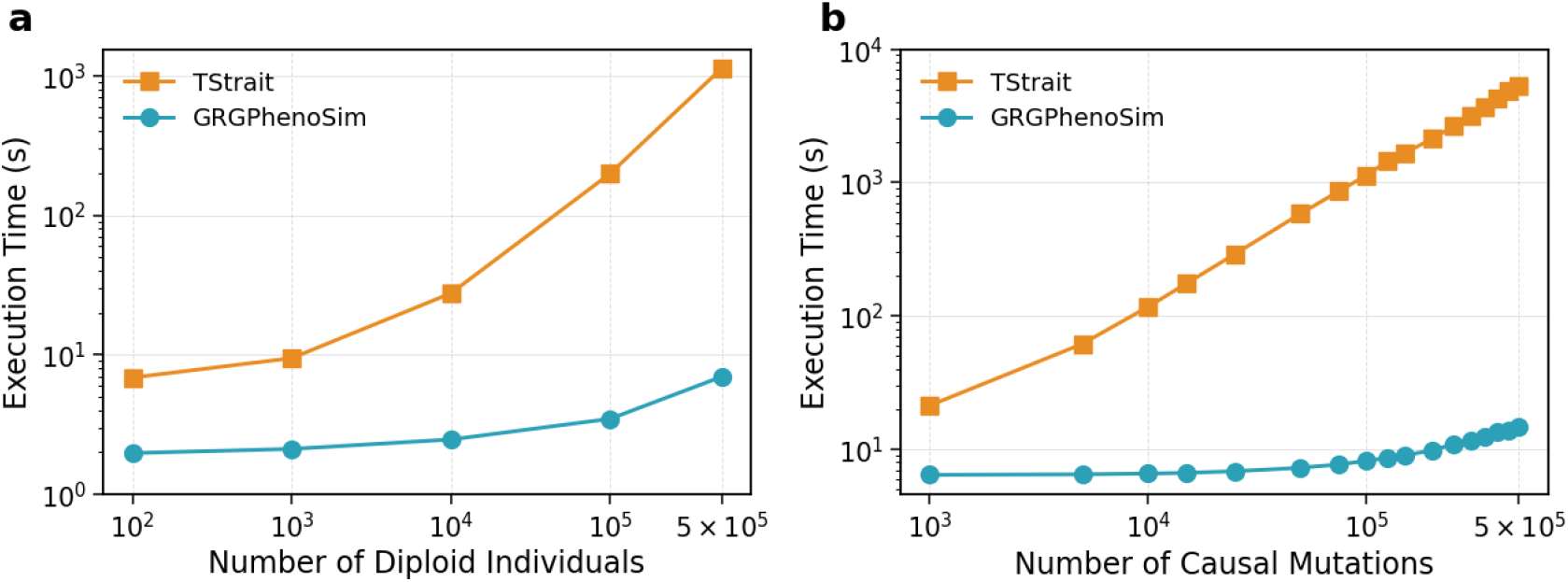
Runtime efficiency tests. The two panels show runtime scaling with (a) increasing sample size and (b) number of causal mutations, illustrating the widening gap between *Grg-PhenoSim* (in orange circles) and *tstrait* (in blue squares) runtimes.

The tests were carried out using a single thread on a compute node with x86 64 architecture, powered by an AMD EPYC 7763 64-Core Processor. A simulated dataset was used to create the GRGs used in the runtime experiments. We used msprime [Baumdicker et al., 2022] to simulate 100 Mbp sequences for up to 500,000 diploid individuals (1,000,000 haploid samples) with a human-like parameter setting (effective population size *N*_*e*_ = 10^4^, recombination rate 10^−8^*/*(bp *×* generation), mutation rate 10^−8^*/*(bp *×* generation)) [DeHaas et al., 2025]. The GRGs were constructed from simulated VCFs. However, the corresponding tree sequences compared against the GRGs were simulated tree sequences.

#### 3.2.1 Scaling with Sample Size

For this experiment, GRGs with 100, 1000, 10,000, 100,000, and 500,000 diploid individuals were used. The corresponding tree sequences were used to run the experiments for *tstrait* [Tagami et al., 2024]. We used 100,000 causal mutations across all the tests.

Overall, we observed a gradual runtime scaling for *GrgPhenoSim* and a steeper scaling for *tstrait* [Tagami et al., 2024], as shown in exhibit (a) in (Fig. 3). Further, *GrgPhenoSim* was significantly faster regardless of the sample size used, with the gap in runtime widening as the sample size increased. At 500,000 individuals, *GrgPhenoSim* was 162x faster than *tstrait* [Tagami et al., 2024].

#### 3.2.2 Scaling with Number of Causal Mutations

For this experiment, the constructed GRG and the corresponding tree sequence for 500,000 diploid individuals were used, with a total of 575,054 mutations in the dataset. The number of causal mutations ranges from 1000 to 500,000.

Once again, we observed a gradual runtime scaling for *GrgPhenoSim* and a steeper scaling for *tstrait* [Tagami et al., 2024], as shown in (Fig. 3b). Moreover, *GrgPhenoSim* was significantly faster for every dataset, and the run-time gap widened as the number of causal mutations increased. At 500,000 causal mutations, *GrgPhenoSim* was 587 times faster than *tstrait* [Tagami et al., 2024].

## 4 Discussion

*GrgPhenoSim* provides a flexible and efficient interface to simulate phenotypes using genotype representation graphs. Taking advantage of the computational speed up from the GRG data structure and wrapping it into simple Python functions, it provides users with the ability to efficiently simulate phenotypes without having to understand the GRG data structure. While *GrgPhenoSim* is faster in all tested settings, the run time advantage becomes even more significant when the sample size is large, which is when repeated phenotype simulations are costly with existing methods. Together with the GWAS function in GRGL and GRG *dot product, GrgPhenoSim* could facilitate the development of statistical genetic methods for biobank-scale datasets, enabling a wider range of GWAS-related applications using GRG functionalities. Last but not least, we demonstrated that GRG dot product can be used to speedup calculations involving standardized genotype matrices, which could facilitate future GRG-based statistical genetics.

## Acknowledgments

This work was partially supported by NIH R35-GM150579 and NSF IIS-2435801. Aditya Syam was partially supported by the Nexus Scholars Program from the College of Arts and Sciences, Cornell University. The authors thank Drew DeHaas, Ziqing Pan, and Richard Border for their valuable input.

